# Numerical analysis of the size-based, shear-induced separation of circulating tumour cells from white blood cells in liquid biopsies

**DOI:** 10.1101/2025.10.13.681808

**Authors:** Benjamin Owen, Timm Krüger, Roslyn Hay, Chiara Ghera, Celine Macaraniag, Jian Zhou, Ian Papautsky

## Abstract

Circulating tumour cells (CTCs) are promising biomarkers for early cancer detection, yet their extreme rarity in blood necessitates efficient separation from white blood cells (WBCs) in lysed liquid biopsies. Inertial microfluidics offers a high-throughput, label-free approach to this challenge by leveraging size-dependent lateral migration. However, experimental observations reveal that WBCs migrate more rapidly than predicted, reducing separation performance. Using 3D lattice-Boltzmann-immersed-boundary-finite-element simulations, we characterized the migration dynamics of CTCs and WBCs in a straight microchannel. Our results reveal that the presence of a CTC enhances WBC cross-streamline migration, providing a mechanistic explanation for WBC contamination in CTC-enriched outlets. The numerical model capturing heterochiral orbital dynamics was validated experimentally, confirming the role of intercellular hydrodynamic interactions. These findings underscore the critical role of intercellular interactions in inertial microfluidic systems and provide guidance for optimizing suspension concentration and channel geometry to improve purity in rare cell isolation.

## 1 Introduction

Cancer remains one of the leading causes of mortality worldwide, accounting for one in six deaths each year [1]. Early diagnosis is crucial to improving survival outcomes. Screening programs for high-risk demographics have been successful in increasing early diagnosis. However, the heterogeneity of cancer types, origin, and clinical presentation limits the feasibility of screening for multiple cancer types, particularly in low-risk demographics. There is a critical need for a low-cost diagnostic test capable of detecting cancer during its early asymptomatic stages of development. One promising strategy involves the identification of circulating tumour cells (CTCs), which are shed into the blood-stream from primary tumours, and can be identified in peripheral blood as early as stage 1 of cancer progression [2]. A major limitation of CTC-based diagnostics is their extreme rarity, around 10–100 CTCs per ml of blood. Therefore, the development of a high-throughput, sensitive detection method is essential to enable reliable identification of CTCs and facilitate early cancer diagnosis.

Inertial microfluidics (IMF) has emerged as a promising label-free technique that exploits fluid inertial in microchannels to manipulate suspended cells through the Segré-Silberberg effect, which drives the lateral migration of particles to equilibrium positions in the channel cross-section [3, 4, 5]. In IMF systems, particles experience inertial lift and drag forces, which generally result in lateral migration and the formation of hydrodynamically bound particle trains [6, 7, 8]. Since migration dynamics depend on the interplay of the channel geometry, the suspending fluid properties, and the suspended particle characteristics [9], IMF offers a passive microfluidic method that does not rely on external fields of active microfluidics [10]. In particular, IMF has been applied for the separation of various particle types, such as blood cells [11], CTCs [12], and bacteria [13].

In straight microchannels, inertial migration generally occurs in two stages while the particles move downstream along the channel axis [14]. The first stage is a fast migration approximately in the radial direction; the second stage is a slower circumferential migration along a heteroclinic orbit (a trajectory connecting unstable and stable equilibrium positions) [14]. While the shape of the heteroclinic orbit is determined by the channel geometry, particle properties (such as size and deformability), and flow conditions (such as *Re*), the location where a particle joins the heteroclinic orbit depends on its initial position.

Particle separation in straight channels can be achieved through two primary mechanisms: 1) particles of different size generally occupy distinct equilibrium positions [15], and 2) they migrate at different lateral velocities [16], reaching the same region of the cross-section at different times. By optimizing channel length and outlet configuration, distinct particle types can be directed into separate outlets [17]. However, in non-dilute suspensions, hydrodynamic interactions between particles can disrupt these trajectories, reducing the purity of the separated streams.

Fig. 1a illustrates the microfluidic device developed by Zhou *et al*. [12], in which blood samples are introduced through the outer inlets, while a cell-free buffer enters through the central inlet. To enhance device performance, red blood cells (RBCs) are removed through lysis, leaving a suspension of white blood cells (WBCs) and CTCs. Due to size-dependent inertial migration, the larger CTCs migrate more rapidly towards the channel center, where stable equilibrium positions are located. The channel length is designed such that only CTCs reach the centerline, enabling their separation from WBCs. However, despite extensive optimization, isolating CTCs from blood remains challenging, as some WBCs also migrate into the focused CTC stream, compromising separation efficiency and purity [18]. This study demonstrates that hydrodynamic interactions between CTCs and WBCs contribute to unintended lateral migration, reducing separation efficiency.

**Figure 1:**
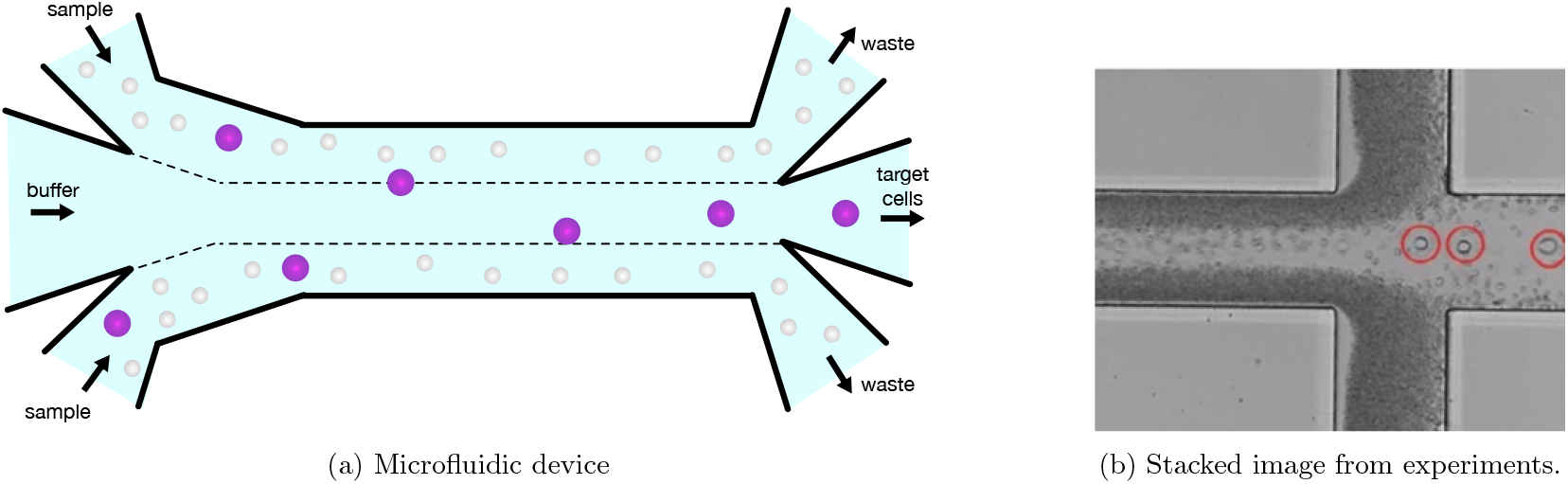
(a) Schematic of the inertial microfluidic device developed by Zhou *et al*. [12], illustrating the lateral migration of CTCs (purple) to the channel centre and their exit through the central outlet. In contrast, WBCs (white) remain primarily in the outer regions of the channel and are directed through the outer outlets. (b) Representative stacked image from Macaraniag *et al*. [18] showing CTCs (circled) exiting through the central outlet. While the majority of WBCs follow the expected path to the outer outlets, a subset is also observed exiting through the central outlet, contrary to theoretical predictions, highlighting the challenge of achieving high-purity CTC isolation.

In this work, we aim to understand the influence of CTCs on the migration dynamics of WBCs and to characterize the velocity-dependent separation behaviour of both cell types. Following a description of the physical (Section 2.1) and numerical (Section 2.2) models, we first analyze the heteroclinic orbits (Section 3.1) and migration velocities (Section 3.2) of individual particles. These modeling results are validated through experiments using beads as CTC analogs. We then examine suspensions containing CTCs and WBCs to assess the role of intercellular interactions in lateral migration, using WBC only suspensions as a reference. We focus on both the macroscopic (Section 3.3) and microscopic (Section 3.4) suspension behavior, providing qualitative and quantitative evidence for the interaction effect. We conclude by discussing the implications of our findings for experimentalists (Section 3.5).

## 2 Methods

### 2.1 Physical model and dimensionless groups

We consider the scenario illustrated in Fig. 1a where the two outer inlets contain a blood sample with WBCs and CTCs, and the inner inlet contains a buffer fluid. Both the suspending fluid of the blood sample and the buffer fluid have the same properties (incompressible, Newtonian, density *ρ*, kinematic viscosity *ν*) and are miscible (no surface tension effects). WBCs and CTCs are treated as resolved neutrally buoyant, spherical objects with diameters *d*_WBC_ and *d*_CTC_, respectively, coupled to the fluid dynamics and interacting with each other hydrodynamically. The no-slip condition holds at the surface of the particles and at the inner surface of the microfluidic device. The numbers of WBCs and CTCs in a simulation are denoted *N*_WBC_ and *N*_CTC_, respectively.

The actual domain of interest is the straight segment of rectangular cross-section between the inlets and outlets as illustrated in Fig. 2c, simply denoted as ‘channel’ with length *ℓ* and cross-sectional height *H* and width *W*. In physical units, the channel dimensions are *H* = 50 *µ*m, *W* = 150 *µ*m and *ℓ* = 30 mm, mimicking the dimensions in [12]. In order to make simulations tractable, we simulate a channel segment of length *L* = *ℓ/*80 and use flow-wise periodic boundary conditions. Fluid and particles leaving the simulation domain at the downstream boundary re-enter the domain at the upstream boundary, and the flow-wise (axial) positions of the particles are tracked over time. This simulation strategy is similar to using a frame co-moving with the particles as they traverse the channel from the inlets to the outlets. The origin of the coordinate system is located at one of the corners of the cross-section, thus the cross-section is defined by 0 *≤ y ≤ W* and 0 *≤z ≤H*. We assume that unperturbed streamlines (*i*.*e*., in the absence of particles) from within the channel continue straight into the three outlets, hence a particle would leave in outlets 1, 2 or 3 when its lateral position is 0 *< y < W/*3, *W/*3 *< y <* 2*W/*3 or 2*W/*3 *< y < W* at the end of the channel, *x* = *ℓ*, respectively.

**Figure 2:**
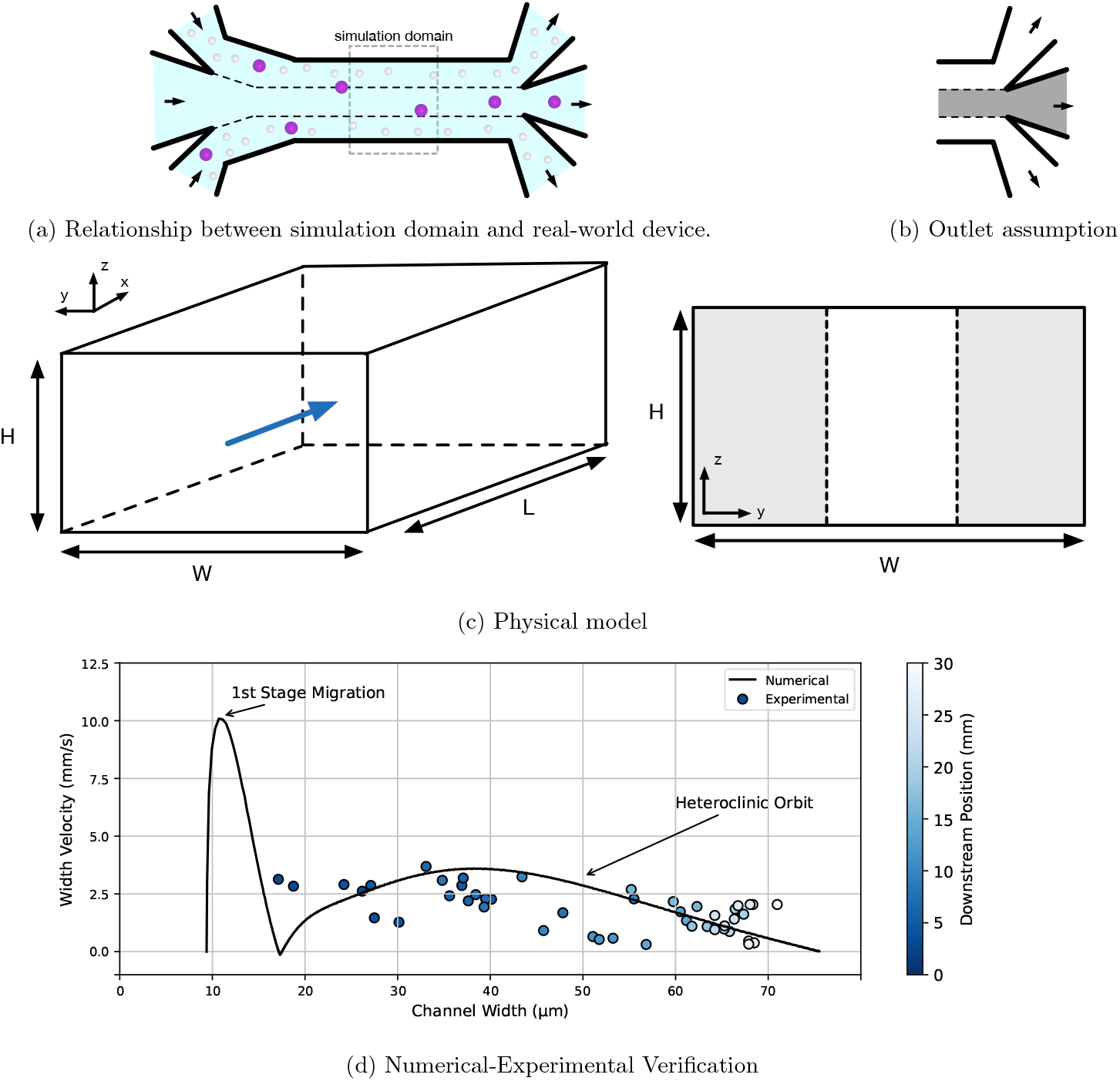
(a) Schematic indicating the simulation domain in relation to the real-world device. (b) Outlet assumptions: particles that migrate to the inner third (grey) are assumed to exit the device through the middle outlet. (c) Schematic of the simulation domain with the blue arrow indicating the axial flow direction. Grey areas in the cross-section indicate the initial locations that white blood cells can be placed. (d) Numerical width velocity predictions are compared to experimental measurements for *χ* = 0.30 particles.

The relevant non-dimensional groups are the channel Reynolds number

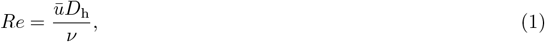

where *ū* is the average velocity in the channel and *D*_h_ = 2*WH/*(*W* + *H*) is the hydraulic diameter of the channel, the confinement for each particle type,

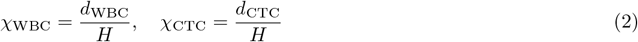

and the channel aspect ratio

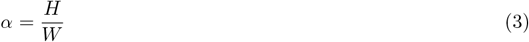

which is *α* = 1*/*3 in the present work.

### 2.2 Numerical model

We employ an in-house solver to simulate the physical system described in Section 2.1. The lattice Boltzmann (LB), finite element (FE) and immersed boundary (IB) methods are used to solve the fluid dynamics, particle dynamics and fluid-particle interaction, respectively. The code is written in C++ and parallelised with the Message Passing Interface. We use the same algorithm as in [19], and our code has been tested previously, for instance in [20]. We validate our code through comparison with experimental data for the device proposed by Zhou et al. [12] in Section 2.6.

Our LB solver involves the Bhatnagar-Gross-Krook (BGK) collision operator with relaxation time *τ* [21] and Guo’s forcing scheme [22] on the D3Q19 lattice [23]. The kinematic viscosity of the fluid is related to *τ* according to *ν* = (*τ −* Δ*t/*2)*/*3 where Δ*t* is the time step. The grid spacing is denoted Δ*x*. A constant body force drives the flow along the channel to reach the desired average axial velocity. The half-way bounce-back method [24] recovers the no-slip condition on the inner surface of the channel, and periodic boundary conditions are used along the flow axis.

Particles are modelled as nearly rigid spherical capsules discretised by a surface mesh with *N*_f_ flat triangular elements in such a way that neighbouring surface points have an average distance of around Δ*x* (*N*_f_ = 2420 for the WBCs and 5120 for the CTCs) [19]. Each capsule experiences local shear, area dilation and bending forces with elastic moduli *k*_S_, *k*_*α*_ and *k*_B_, as well as volume and surface conservation forces with moduli *k*_V_ and *k*_A_, respectively [20]. The softness of a particle with radius *a* is determined by the Laplace number

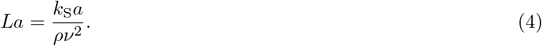

In the current work, particles are simulated in the near-rigid limit with *La* = 100. Additionally, *k*_*α*_ and *k*_B_ are fixed by *k*_*α*_*/k*_S_ = 2 and *k*_B_*/*(*k*_S_*a*^2^) = 2.87 *·* 10^*−*3^, following [19].

The immersed boundary method [25] provides the two-way interaction between the particles and the fluid and recovers the no-slip condition at the surface of the particles. We use Peskin’s three-point stencil for velocity interpolation and force spreading.

Since the suspension is dilute, hydrodynamic forces are sufficient to maintain a fluid layer of at least 2Δ*x* between any two particles and between particles and the walls, hence contact models are unnecessary.

### 2.3 Simulation set-up and parameters

A schematic of the simulation setup is shown in Fig. 2c. The flow is driven along the *x*-axis by a body force in such a way that the desired average velocity *ū* is achieved. The simulated Reynolds number is *Re* = 50 in all cases. All relevant physical and numerical parameters for the geometry, fluid and particles are collected in Tab. 1.

**Table 1:**
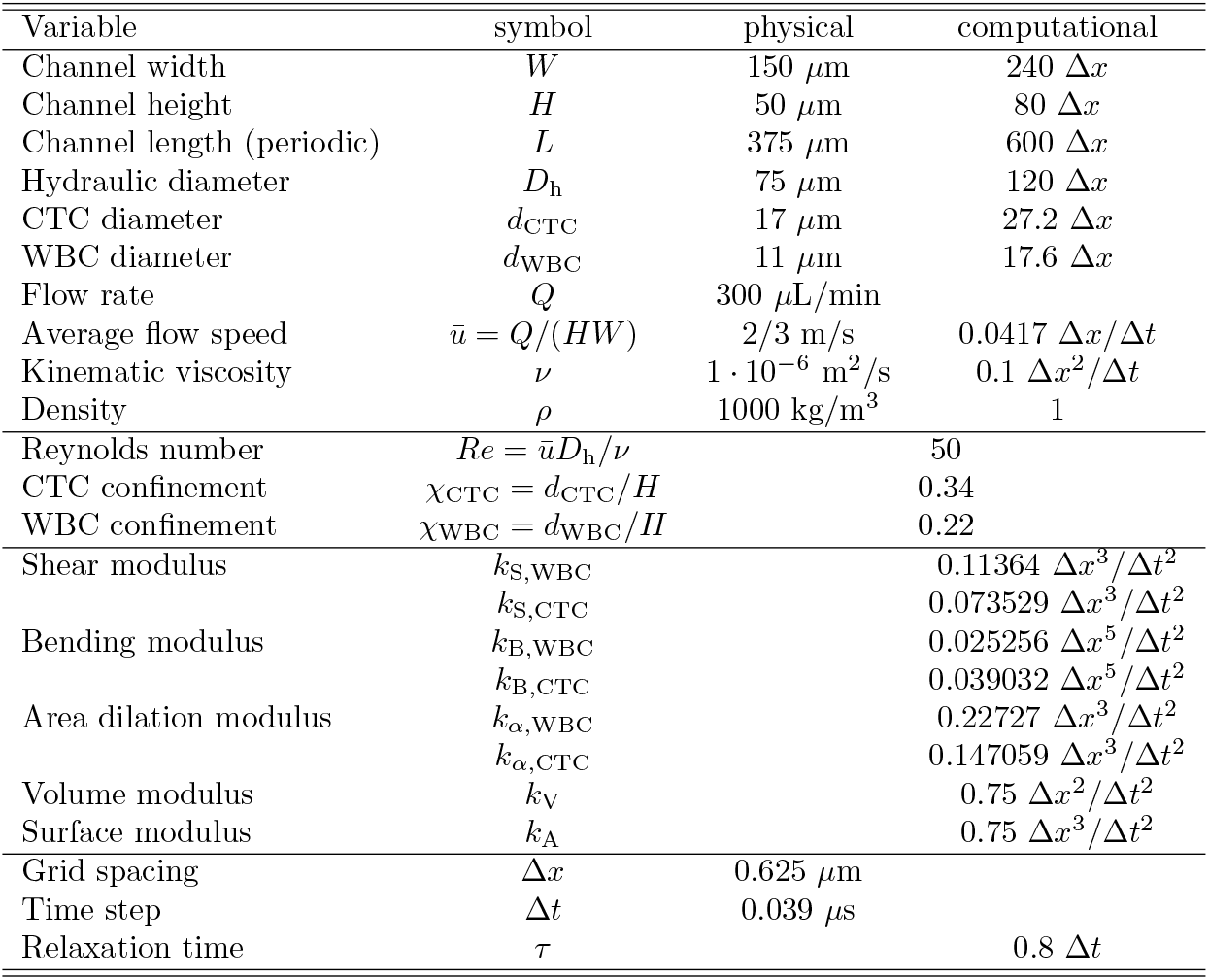
List of physical and numerical simulation parameters.

A specified number of WBCs and CTCs are randomly initialised within the two outer channel segments (*y < W/*3 and *y >* 2*W/*3), thus approximating particles entering the channel through the two blood inlets. The flow is then started, and the simulations run until an abort condition is reached, for example when the CTC reaches the end of the channel at *x* = *ℓ*.

Due to the smaller simulation domain, *L* = *ℓ/*80, we use *N*_CTC_ = 1 and *N*_WBC_ = 8, which results in a volumetric particle concentration of 0.2%, matching experiments in [12]. However, the number ratio of WBCs and CTCs in experiments [12] is around ???, thus our simulation results need to be interpreted accordingly.

The observables of interest are the cross-sectional positions, (*y, z*), and migration velocities, (*v*_*y*_, *v*_*z*_), of all particles as function of time, *t*, and axial particle position, *x*. Particle positions are normalised by the dimensions of the channel cross-section, and particle velocities are normalised by the average flow velocity, *ū*.

### 2.4 Microfluidic device fabrication

The microfluidic device consisted of a straight rectangular microchannel with dimensions of 150 *µ*m (width) × 50 *µ*m (height) × 30 mm (length), incorporating three inlets—two for sample introduction and one central inlet for buffer—and three outlets. The central outlet was designed to collect larger particles, while the two outer outlets collected smaller ones. Devices were fabricated by casting polydimethylsiloxane (PDMS) onto a photoresist master to form the microchannel structures, following established protocols as described previously [26]. Dry photoresist films (ADEX 50, DJ MicroLaminates Inc., Sudbury, MA, USA) were patterned on 75 mm diameter silicon wafers to define the channel geometry. PDMS (Sylgard 184, Dow Corning, Midland, MI, USA) was cast onto the master and cured. The resulting PDMS replicas were then bonded to 25 mm × 75 mm glass slides (Fisher Scientific Inc., Hampton, NH, USA) following 20% oxygen plasma treatment (PE-50, Plasma Etch Inc., Carson City, NV, USA) for 20 s at 300 W with a continuous oxygen flow of 10 cc/min. Inlet and outlet ports were cored using a 1.5 mm diameter biopsy punch (TedPella Inc., Redding, CA, USA) prior to bonding.

### 2.5 Microfluidic experiments and data analysis

Fluorescent polystyrene beads (15.45 *µ*m diameter; Life Technologies Inc.) were used to mimic CTCs. Beads were suspended in a saline solution containing 0.9% (v/v) NaCl and 0.2% (v/v) Tween-20 (Fisher Scientific Inc.) to prevent aggregation and minimize clogging. Suspensions were prepared at volume fractions of 1% and 0.1% (v/v), and introduced into the device via PTFE tubing (Cole-Parmer) connected to a syringe pump (Legato 200, KD Scientific Inc., Holliston, MA, USA). A total flow rate of 300 *µ*L/min was used, corresponding to *Re* = 50.

Particle migration was recorded using a high-speed camera (Mini AX200, Photron USA Inc., San Diego, CA, USA) at 10,000 fps with a shutter speed of 2.5 *µ*s on an inverted microscope (IX83, Olympus America, Center Valley, PA, USA). Raw image sequences were converted to TIFF format and analyzed using ImageJ^®^. Individual particle trajectories were isolated using the “Make Substack” tool, followed by thresholding and analysis with the “Analyze Particles” function.

Lateral migration velocities were measured for five individual particles at each downstream position, beginning at 1 mm and continuing at 2 mm intervals along the remaining channel length. The migration velocity *V*_*m*_ was calculated using the formula:

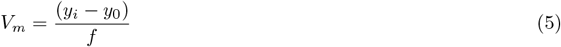

where *y*_*i*_ is the lateral position at frame *i, y*_0_ is the initial position, and *f* is the frame rate. Measurements were taken every five frames.

### 2.6 Validation

The validity of the simulation results was confirmed through comparison with experimental data. Specifically, we compared the lateral velocity of particles (*χ* = 0.30) in the channel width direction measured experimentally with predictions from the numerical model. The simulations were performed using a single isolated particle. In contrast, capturing the behaviour of an individual particle experimentally is impractical. However, the suspension is sufficiently dilute (0.01%) to minimise particle-particle interactions, ensuring that the simulation conditions were representative of the experimental environment.

Fig. 2d shows a comparison between numerically predicted and experimentally measured lateral velocities during particle migration across the channel width. Experimental measurements were taken at various axial positions, as indicated by the colour gradient. The numerical results exhibit two distinct velocity peaks. The larger peak, located near the channel wall, corresponds to the first stage of migration, from the initial position to the heteroclinic orbit. This stage is dependent on the initial position, and due to the difficulty in measuring initial positions in the width and depth directions simultaneously, robust comparison of numerical and experimental measurements for this stage of migration is not feasible. We therefore exclude any experimental measurements made before *x* = 2 mm, allowing for direct comparison of the migration behaviour associated with the heteroclinic orbit, represented by the smaller peak in Fig. 2d. A good agreement is observed between the experimental and numerical measurements, both exhibiting the matching parabolic profiles.

## 3 Results

We first present the migration trajectories of single particles, defining the shape of the heteroclinic orbits, as a function of confinement (Section 3.1) before discussing the migration velocity of a single particle along the heteroclinic orbit (Section 3.2). The macroscopic migration behaviour of CTCs and WBCs in dilute suspensions is investigated in Section 3.3, followed by an analysis of the microscopic interactions of CTCs and WBCs and its influence of the WBC dynamics (Section 3.4). We conclude by discussing the implications of our findings for experimentalists (Section 3.5).

### 3.1 Migration of single particles and shape of heteroclinic orbits

To establish a baseline for identifying particle-particle effects, we investigate the migration behaviour of single particles with different confinement values. Particles are simulated in the near-rigid limit with *La* = 100, and particle confinement is varied within the range of *χ ∈* [0.18, 0.38]. For reference, in the real-world geometry used by Zhou *et al*. [12], a WBC corresponds to *χ*_WBC_ = 0.22, while a CTC corresponds to *χ*_CTC_ = 0.34.

Fig. 3a and Fig. 3b show the lateral migration of particles of size *χ* = 0.22 and *χ* = 0.34, respectively. Each particle is initialised at various locations, identical for both particle sizes. Both small and large particles exhibit an initial rapid radial migration, followed by a slower migration along the heteroclinic orbit. The characteristic shape of these orbits is evident in the overlapping segments of the particle trajectories. However, the location at which a particle transitions onto the heteroclinic orbit is strongly dependent on its initial position. Note that particles do not necessarily follow the shortest possible path from the initial position to the heteroclinic orbit. Particles initialised near the channel wall are primarily dominated by the wall repulsion force during the first stage of migration. In contrast, particles starting closer to the channel centre experience dominant shear gradient lift force, which drives them toward regions of higher shear before aligning with the heteroclinic orbit.

**Figure 3:**
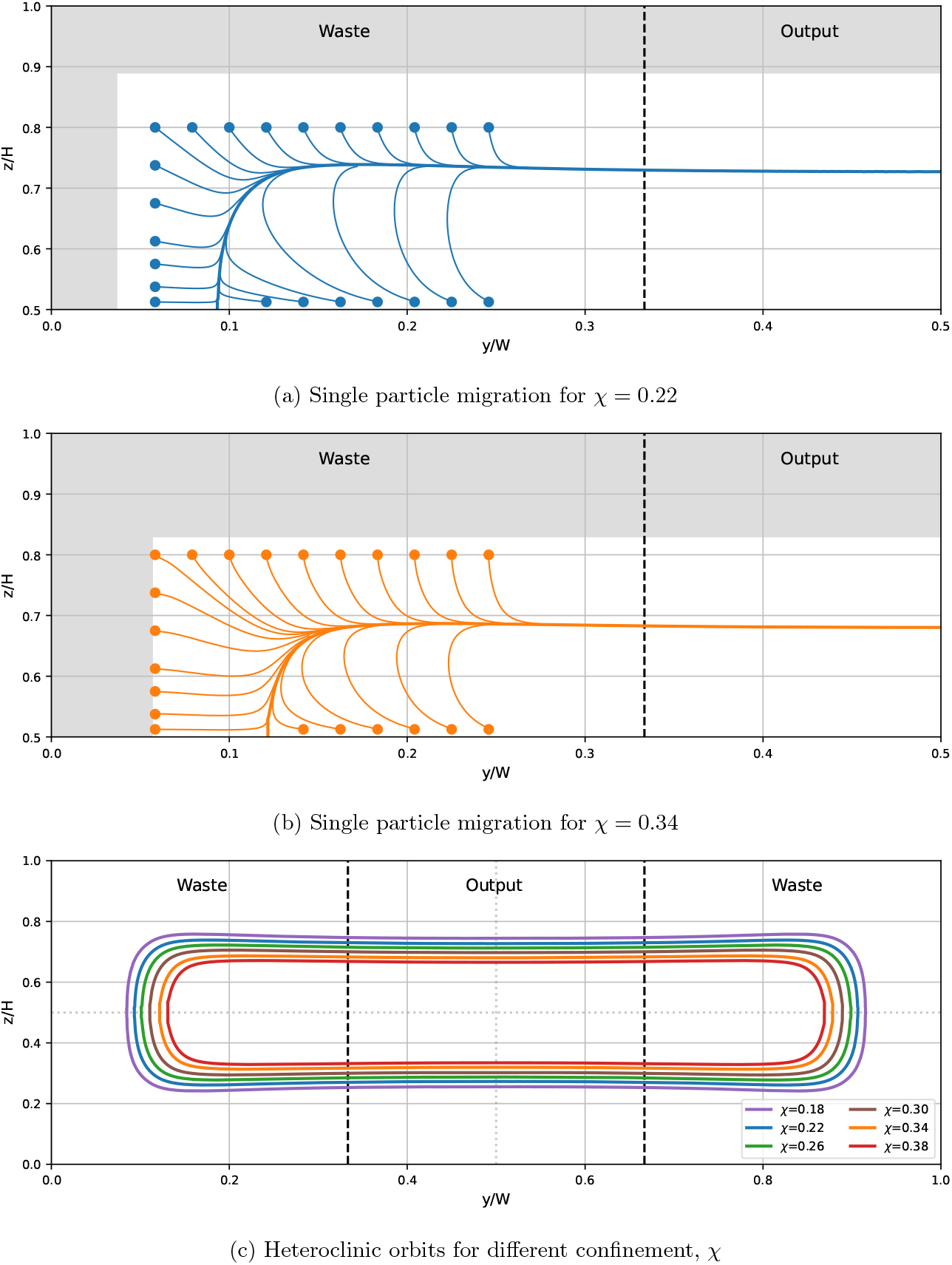
Cross-sectional view of lateral migration path for a single particle with (a) *χ* = 0.22 and (b) *χ* = 0.34 when initialised at different positions. Initial positions are denoted by a circle, while the equilibrium position is denoted by a square. Due to symmetry of the rectangular channel cross-section, only one quarter of the cross-section is considered. Each line denotes the path of the particle centre over the channel cross-section. Grey areas denote the locations where the particle centre of mass cannot exist due to the finite size of the particle and the presence of the channel wall. (c) Total cross-sectional view of heteroclinic orbits for a single particle with confinement values in the range *χ ∈* [0.18, 0.38]. The vertical dashed lines corresponds to *y* = *W/*3 and *y* = 2*W/*3, the dividing lines between the portions of the flow entering the first, second and third outlet, respectively.

Fig. 3c reveals the dependency of the shape of the heteroclinic orbit on the particle confinement. Larger particles result in heteroclinic orbits located closer to the channel centre, which is due to disproportionally stronger wall repulsion forces for larger particles

### 3.2 Migration velocity along heteroclinic orbits

Particles introduced through either of the two suspension inlets require time to migrate laterally from these sample streams near the channel sidewalls toward the central output stream, where equilibrium positions are located. If the channel length is insufficient for complete lateral migration, the final distribution of particles across the outlet stream depends on their migration velocity. In such cases, size-based separation can still occur—not due to differences in equilibrium positions, but as a consequence of differential migration velocities between particles of varying size.

Fig. 4a–d shows the lateral position, *y*, of a single particle for different confinement values and four different initial locations as a function of the downstream (axial) position, *x*, of the particle. These four initial positions representative of the extreme locations of the particles in the outer third of the channel and are depicted in the inset of Fig. 4e. We observe that larger particles generally migrate faster along the width direction. Specifically, the downstream position *x*_cr_ at which a particle reaches *y*_cr_ = *W/*3 and transitions from the waste stream to the output stream occurs earlier for larger particles than for smaller ones.

**Figure 4:**
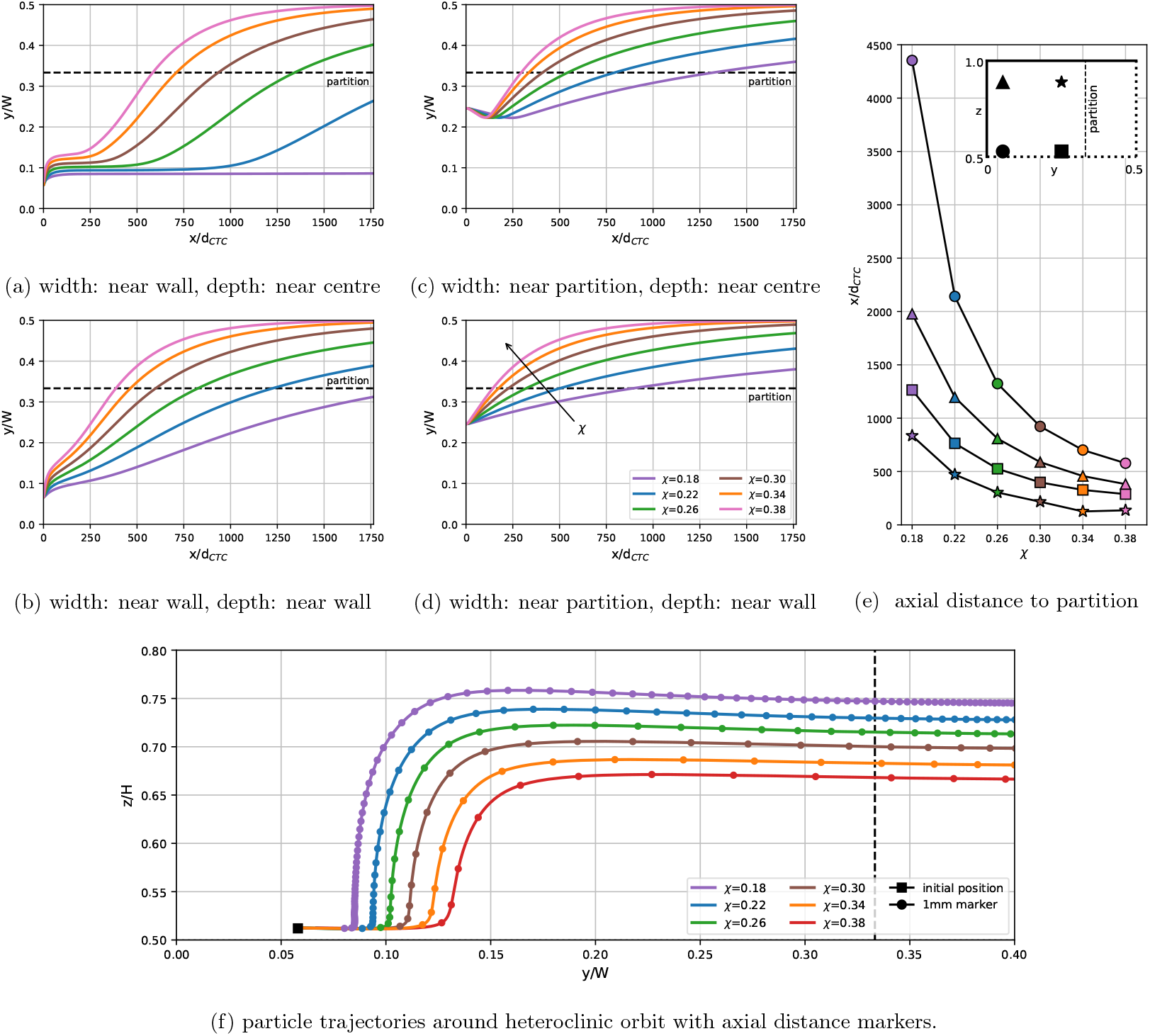
(a–d) Particle migration in the width direction as the particle flows in the axial direction. Particle confinement is varied. Two initial positions in the width direction, close to wall and close to partition, and two initial positions in the depth direction, close to centre and close to wall, are considered. The dotted line indicates where the particles have migrated sufficiently to cross the partition between outlets. The axial distance range shown in the figure is indicative of a real-world device, approximately 3 cm long assuming a CTC diameter of 17 *µ*m. (e) Axial distance travelled for particles of different confinement from each of the four initial positions indicated by different colours corresponding with the schematic (inset). (f) Cross-sectional particle migration for different confinement, *χ*. Markers indicate an axial distance of 1 mm.

Fig. 4e summarises the data from Fig. 4a–d in terms of the axial distance *x*_cr_ required for a particle to reach *y*_cr_. Regions with a large gradient magnitude |d*x*_cr_*/*d*χ*| are of particular interest, as they indicate that particles with small size differences require appreciably different migration times to reach the output stream, thus enabling velocity-based separation. The gradient is more pronounced for particles initially located closer to the sidewall, suggesting that separation performance may be enhanced by controlling the inflow locations of the particles.

Our analysis indicates that the majority of the time a particle requires to reach *y*_cr_ is spent migrating along the heteroclinic orbit (data not shown). Particles initially close to *y*_cr_ spend about half of their migration time on the heteroclinic orbit, while particles starting near the side wall spend up to 90% of their time on the heteroclinic orbit.

Therefore, the time to cross-over is largely determined by the location at which a particle joins the heteroclinic orbit. Fig. 4f presents a modified version of Fig. 3c with markers added to the heteroclinic orbits to indicate the axial distance required to migrate along the orbit. We found that the migration velocity on the heteroclinic orbit is only a function of current position and does not depend on the history of the particle (see Fig. A1). A particle migrating from one marker to the next requires a constant axial distance *δx* where *δx* = 1 mm for a CTC in the device used in [12]. Lateral migration is generally slow in the vertical segment of the heteroclinic orbit (*y ≈* 0.1*W*) and near the equilibrium position (*y ≈ W/*2). Particles migrate fastest in the intermediate region between *y* = 0.2*W* and *y* = 0.3*W*. The slow migration around *y ≈* 0.1*W* is explained by the presence of an unstable equilibrium position on the *y*-axis at *z* = *H/*2 while the slow migration toward *y* = *W/*2 is caused by the presence of the equilibrium position at *y* = *W/*2. In the region *y ≈* 0.1*W*, the lateral migration velocity exhibits strong size dependence, with larger particles migrating much faster. This observation suggests an additional mechanism for velocity-dependent separation by controlling particle entry locations within the channel.

### 3.3 Macroscopic WBC migration with and without CTC present

We investigate the macroscopic characteristics of WBC migration in a dilute suspension (*χ*_WBC_ = 0.22) with or without a single CTC (*χ*_CTC_ = 0.34) under the same flow conditions as before (*Re* = 50). Eight WBCs are included in each simulation, with volume fraction of 0.2%. We consider three different scenarios: 1) no CTC present; 2) one CTC present, originally located outside its heteroclinic orbit; 3) one CTC present, originally located inside its heteroclinic orbit. For each scenario, we simulated 15 different realizations with different random initial positions for the WBCs but the same initial position for the CTC. All particles are initially located in the two sample inlet regions (*y*_0_ *< W/*3 and *y*_0_ *>* 2*W/*3).

Fig. 5 summarizes the results of all three scenarios, with S1 shown in the left column, S2 in the middle, S3 in the right. Panels (a–c) in the first row show stacks of top view (*x*-*y*) trajectories, where CTC trajectories are highlighted in red, and the WBC trajectories in grey. In the absence of a CTC (Fig. 5a), WBCs travel downstream for over 1000 CTC diameters before reaching the channel centerline. In contrast, when CTC is present (Fig. 5b and Fig. 5c), WBCs reach the centerline within approximately 500 CTC diameters. Additionally, WBC trajectories exhibit increased fluctuation in the presence of a CTC. These observations suggest that the presence of a CTC accelerates the lateral migration of some WBCs toward the channel centre, therefore reducing the purity of CTCs in the output stream. Importantly, the overall shape of the WBC trajectories appears to be mirror symmetric with respect to the centreline, although the CTCs are only located in the region *y > W/*2, implying that the CTCs affect the WBCs on both sides of the channel.

**Figure 5:**
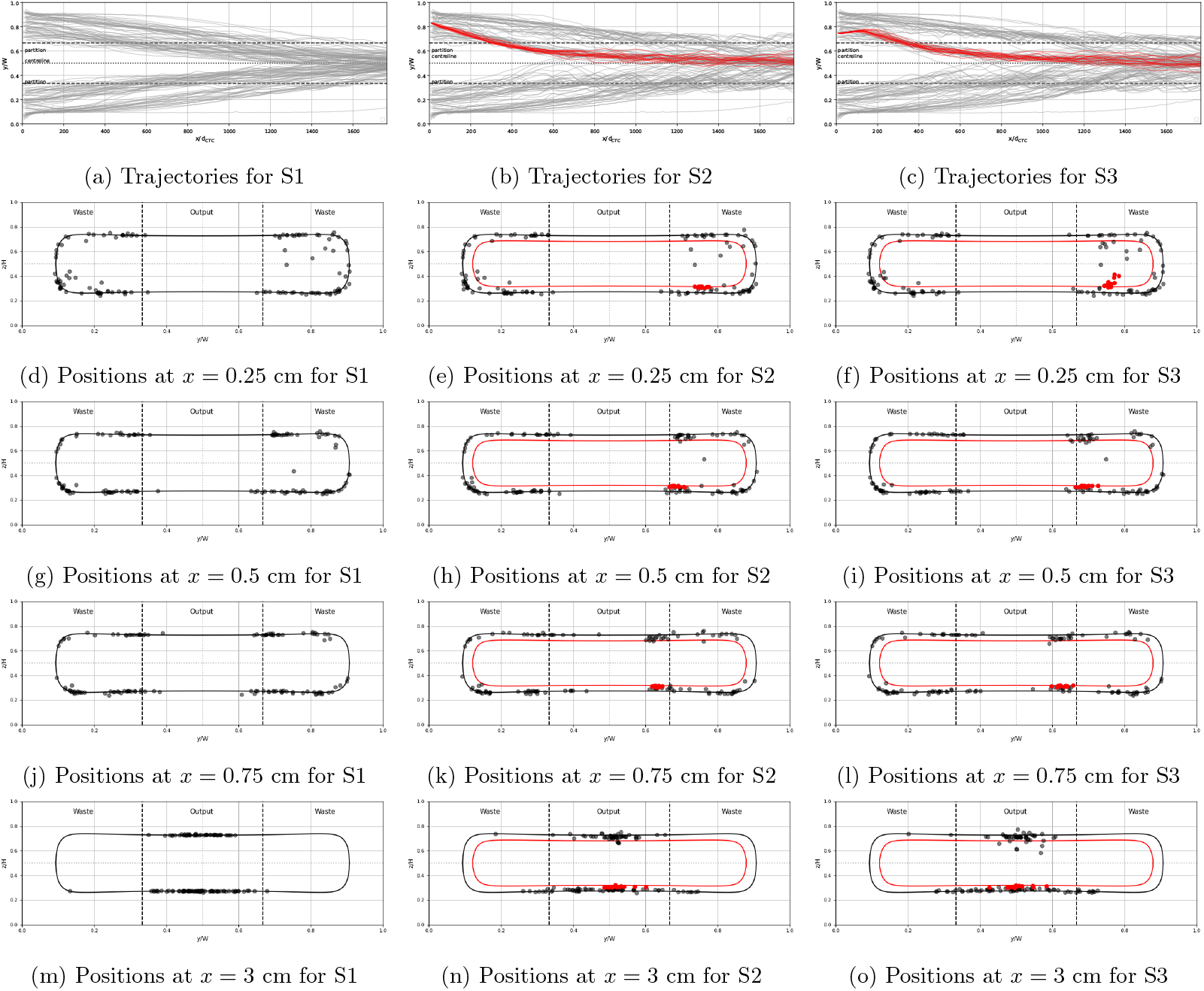
Particle migration behaviour for three scenarios: 1) no CTC present (S1, left column), 2) single CTC initially outside its heteroclinic orbit (S2, middle column) and 3) single CTC initially inside its heteroclinic orbit (S3, right column). (a–c) Stack of trajectories (top view) of eight WBCs from 15 simulations. CTC trajectories are shown in black. WBC trajectories are coloured by initial quadrant location with respect to the initial position of the CTC. All trajectories are shown from their initial positions up to an axial location of *x* = 3 cm. (d–f) Cross-sectional positions of WBCs (black) and CTCs (red) at axial location *x* = 0.25 cm. The corresponding heteroclinic orbits for both particle types are shown as thin lines in corresponding colours. (g–i) The same for *x* = 0.5 cm. (j–l) The same for *x* = 0.75 cm. (m–o) The same for *x* = 3 cm.

The remaining panels of Fig. 5 show the cross-sectional locations of the particles as they flow downstream. Panels (d–f) display the particle positions at *x* = 0.25 cm, panels (g–i) at *x* = 0.5 cm, panels (j–l) at *x* = 0.75 cm, and panels (m–o) at *x* = 3 cm. In scenario 1 (left column), in the absence of a CTC, the WBCs behave similarly to isolated WBCs by first reaching their heteroclinic orbit and then migrating toward the output channel. While WBCs are not always precisely located on the heteroclinic orbit, this deviation is likely ascribed to hydrodynamic interactions between the WBCs.

In scenarios 2 and 3 (middle and right columns), where CTC is present, the CTCs rapidly reach the heteroclinic orbit and then migrate along it toward the output channel. WBCs exhibit broadly similar behaviour to scenario 1, but with notable differences. First, WBCs tend to deviate more from their heteroclinic orbit, particularly in regions vertically opposite the CTC. This deviation is accompanied by clustering of WBCs on the side opposite the CTC, suggesting the possible formation of staggered pairs between CTCs and WBCs. Second, WBCs on both sides of the channel migrate more rapidly toward the channel centerline when a CTC is present. Overall, no significant differences are observed between scenarios 2 and 3.

The observed differences between scenario 1 on scenarios 2 and 3 appear to result from hydrodynamic interactions between CTCs and WBCs. In particular, the formation of staggered pairs between CTCs and WBCs may contribute to the accelerated migration of WBCs opposite of the CTC. Additionally, WBCs on both sides of the channel exhibit faster migration toward the centerline when a CTC is present, despite the CTC being confined to one side. This suggests that the influence of the CTC extends across the channel, although the mechanism driving the enhanced migration of WBCs on the opposite side remains unresolved.

### 3.4 Microscopic WBC dynamics

To further investigate the mechanism underlying the increased rate of WBC migration in the presence of a CTC, we analyse particle-particle interaction in one of the simulations included in Section 3.3 as shown in Fig. 6a. In this example, a CTC (red) appears to draw a WBC (blue) from the opposite inlet across the channel centerline. Note that an isolated cell would be unable to cross the channel centre due to the symmetry of the fluid velocity profile.

**Figure 6:**
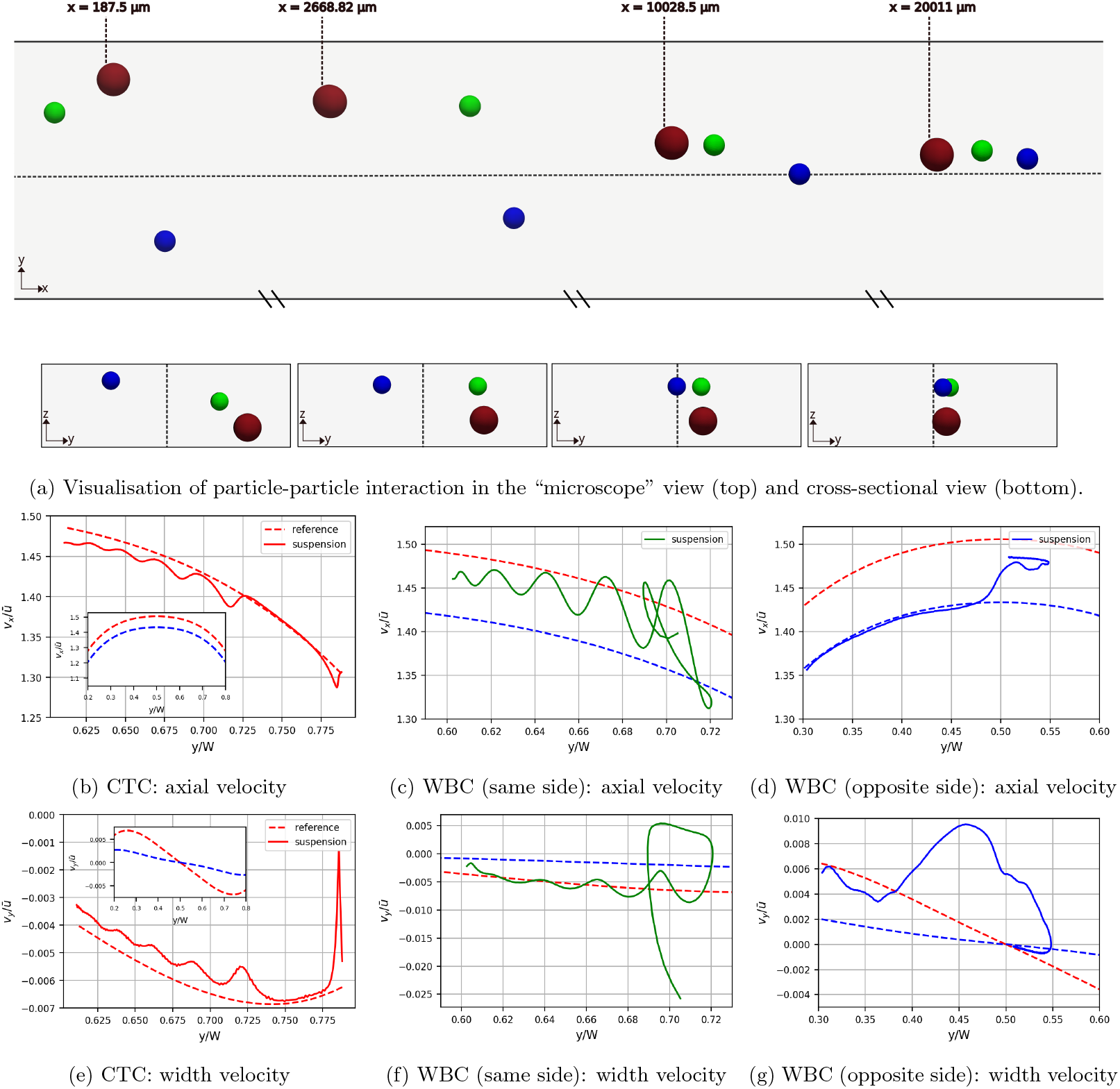
Example particle-particle interaction as cells pass through the channel. Visualisation of the interaction is shown in (a) from top-down or “microscope” view and from the cross-sectional view. A CTC (red) interacts with a WBC initially located on the same side of the channel as the CTC (green) and a WBC initially located on the opposite side of the channel (blue). The axial velocity of each particle during their lateral migration in shown in panels (b) for the CTC, (c) for the same side WBC, and (d) for the opposite side WBC. The lateral velocity of each particle during their lateral migration in shown in panels (e) for the CTC, (f) for the same side WBC, and (g) for the opposite side WBC. Dashed lines show single particle migration behaviours for the CTC (red) and WBC (blue) for comparison in panels (b–g). Full single particle migration behaviours across the full channel width are shown in the inset of (b) for the axial velocity and (e) for the lateral velocity.

The CTC also interacts with another WBC (green), this time from the same inlet as itself, eventually forming a train of cells. Note that in the simulation which this figure visualises, six other WBCs are present but are not shown in order to focus the visualisation since the interaction of other WBCs is minor relative to the WBCs shown.

To quantify the effect of particle-particle interactions, we use the migration velocities of isolated single particles in the lateral direction as a reference. As established in Section 3.2, single particles first migrate to the heteroclinic orbit and then proceed along it. Once on the orbit, particle migration behaviour becomes independent of initial position, allowing prediction of both the axial and lateral velocities at any given position. This predictable heteroclinic orbit behaviour in the absence of neighbouring particles serves as the reference, shown in the inset of Fig. 6b for axial velocity and the inset of Fig. 6e for lateral velocity. We then compare the real velocity at a given lateral position in the suspension case to the expected velocity from the isolated case. To simplify the analysis and reduce the total number of interactions, we used a reduced suspension case in which only the CTC and two WBCs shown in Fig. 6a are present. A train of particles forms in this reduced system in a manner consistent with the interaction visualised in Fig. 6a.

All particles in the suspension showed deviations from the single particle reference curves. For the CTC (red) and the WBC located on the same side of the channel (green), both axial and lateral velocities exhibited oscillations not present in the isolated particle cases. These oscillations are attributed to particle-particle interactions, consistent with findings from previous studies on pairwise dynamics [6, 7, 27].

A clear difference in mean behaviour was observed between the CTC and the WBCs when compared to their representative reference cases. The CTC exhibits reduced axial and lateral velocities relative to an isolated CTC. In contrast, WBCs in suspension migrate more rapidly in both directions, with the increase in lateral velocity being more pronounced than in the axial direction. For the WBC located on the opposite side of the channel (blue), axial velocity closely follows the reference curve while the particle remains on its original side. However, upon crossing the channel centre and entering the region on the same side as the CTC, the axial velocity increases. The lateral velocity of this WBC (blue) remains elevated throughout the migration up to the point of close proximity with the CTC.

These findings demonstrate that the presence of a CTC can increase the migration rate of a WBC, increasing the lateral velocity of a WBC, even when initially located on the opposite side of the channel. In practice this finding means that suspensions must be sufficiently dilute in order to prevent such particle-particle interaction occuring.

To demonstrate how the CTC increases WBC migration rates beyond a single example, we investigate the lateral velocities of particles that are closest to each other, at various time points throughout the simulations included in Fig. 5 (approximately 800 times per simulation). We show the distribution and density of lateral velocities for a given axial distance between particles in the WBC only suspension case (S1) and the CTC suspension cases where the CTC is initially located outside the heteroclinic orbit (S2) and outside the heteroclinic orbit(S3) in Fig. 7. Each plot contains data extracted from all 15 configurations in a similar way to the stacked images of Fig. 5. The maximum lateral velocity for a single WBC and a single CTC are included in Fig. 7 as annotated grey dashed lines for reference. All distributions present in a head-and-tail configuration, with larger lateral velocities at closer proximities, indicating the increased velocity is linked to particle-particle interactions. The lateral velocities regularly exceed the single particle maxima for the CTC and WBC, indicating the effect of the particle-particle interaction has a large influence on the migration behaviour of the particles. We see that in both the CTC cases, the particles are closer together more often, and have larger lateral velocities more often than in the WBC only case. This observation has two consequences. Firstly, any particle-particle interaction increases lateral migration. Secondly, a larger particle will influence a smaller particle more strongly. Both consequences can be negated by ensuring suspensions are sufficiently diluted to decrease the chance of particle-particle interaction. Note the difference between the peak density of axial distances between particles is expected since a CTC and WBC pair will have a smaller equilibrium axial distance than a WBC-WBC pair [28].

**Figure 7:**
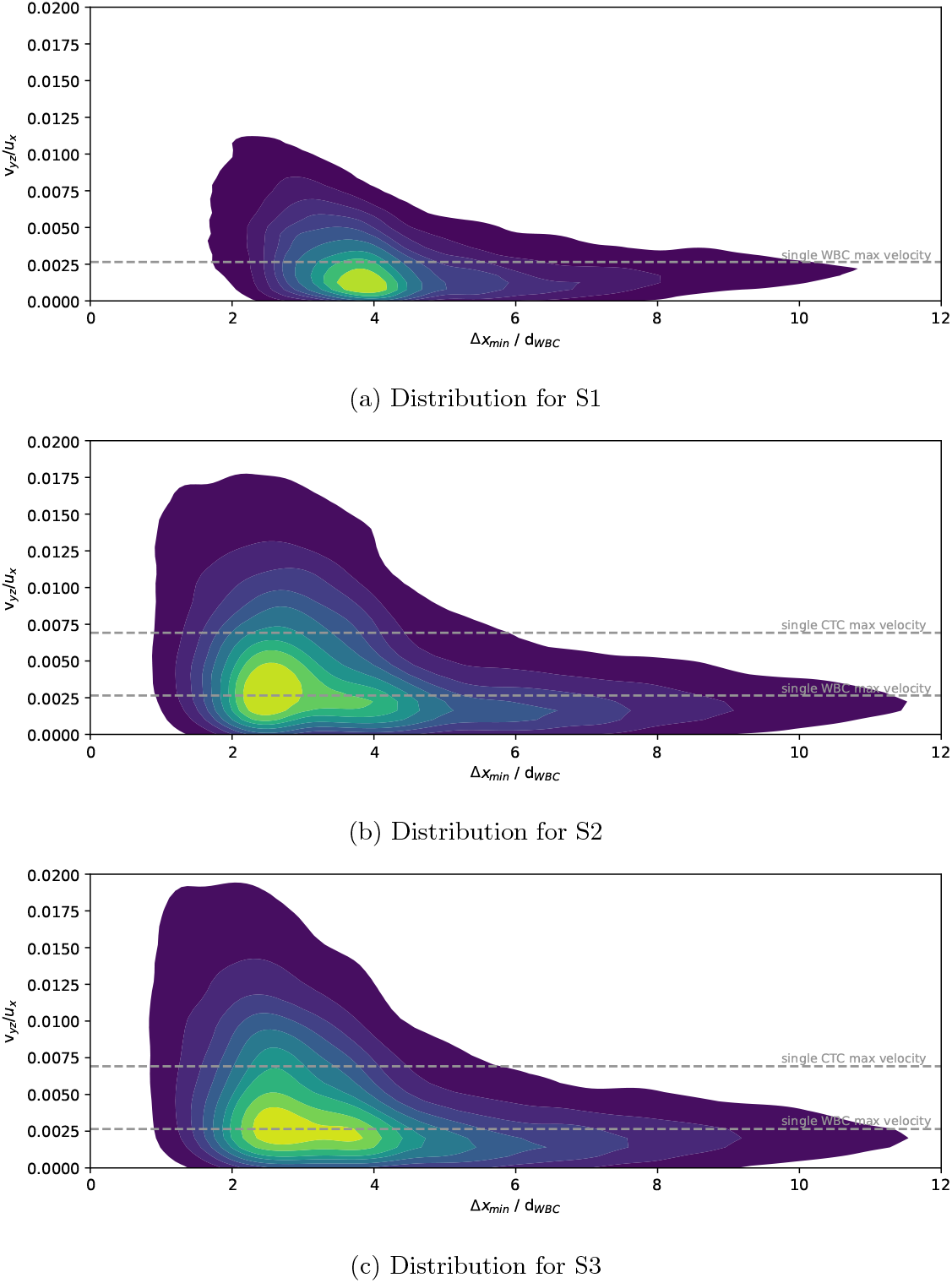
Distribution map of lateral velocity with axial proximity for the nearest neighbours for the three scenarios shown in Fig. 5, (a) no CTC present, (b) single CTC initially outside its heteroclinic orbit and (c) single CTC initially inside its heteroclinic orbit. Each map contains data for the same 15 simulations stacked in Fig. 5.

One assumption we have made in our simulation set-up is using a reduced domain length and periodic boundary conditions. This assumption means a reduced number of WBCs in the domain, however a fast moving particle is able to repeatedly catch-up and overtake other particles with each occurrence counted as a separate interaction. Finally, in Fig. 8 we show that CTCs can interact with a large number of WBCs (as periodic images in the simulation). This finding indicates that a given CTC within the device has multiple opportunities to drag WBCs into the target stream, further demonstrating the need to fully understand the mechanism behind the particle-particle interaction and to ensure the suspension is sufficiently dilute.

**Figure 8:**
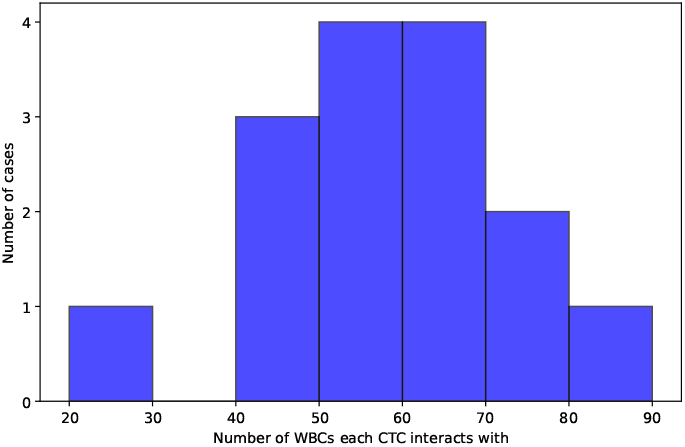
Histogram of the number of WBCs that a CTC has the opportunity to interact with as it travels the axial distance of a representative real-world device (3 cm). 15 simulations from S2 are included.

### 3.5 Implications for experimentalists

The findings of this study have several important implications for the design and optimization of IMF devices intended for separating CTCs from WBCs. A central observation is that particles predominantly migrate along heteroclinic orbits—trajectories that connect unstable to stable equilibrium positions within the channel cross-section. These orbits are strongly size-dependent, with larger particles tending to follow orbits located closer to the channel centerline, while smaller particles remain nearer the channel walls.

Although the migration velocity along a heteroclinic orbit is independent of a particle’s initial position, the initial location determines the segment of the orbit that the particle joins. This dependency has practical implications for device design, as inlet positioning and flow focusing strategies can influence the time required for particles to reach their equilibrium positions. Moreover, the migration velocity is not uniform along the orbit; particles migrate more slowly in the depth direction than in the width direction. This anisotropy implies that channel length requirements may differ depending on the dominant migration direction for a given particle type. The spatial variation in migration velocity also depends on particle size. Differences in migration behaviour between particle sizes are more pronounced when particles enter the heteroclinic orbit near the channel wall in the width direction and near the centre in the depth direction. This spatial sensitivity should be carefully considered when designing channel geometries and selecting flow rates to maximize separation resolution.

Another critical consideration is the role of particle-particle interactions, which can further complicate migration dynamics. While individual particles tend to return to their heteroclinic orbits after perturbations, interactions in non-dilute suspensions can temporarily displace particles from their expected paths. Notably, larger particles can induce faster migration of nearby smaller particles compared to their isolated behavior. This effect can lead to unexpected crossover events and reduced separation purity, as observed in experimental systems where WBCs appear in the CTC outlet.

Our findings highlight a key limitation of relying solely on single-particle theoretical predictions for device design. Such models are only valid when the suspension is sufficiently dilute to render particle-particle interactions negligible. However, no universal criterion currently exists for defining what constitutes a “sufficiently dilute” suspension in the context of IMF. As a result, experimental validation remains essential, and caution should be exercised when extrapolating from idealized models to practical device performance.

Finally, the formation of particle pairs or trains introduces new collective migration behaviors. These emergent heteroclinic orbits depend on the size and configuration of the particles within the pair or train, potentially altering the expected separation dynamics. This finding further emphasizes the need for models that incorporate multi-particle interactions.

Taken together, these insights highlight the importance of accounting for both single-particle dynamics and inter-particle interactions in the IMF design. Experimentalists should consider not only particle size and flow conditions but also initial positioning, channel geometry, and suspension concentration to optimize separation performance. These considerations provide a foundation for refining experimental protocols and improving the reliability of rare cell separation in clinical and research applications.

## 4 Conclusions

In this study, we use an in-house lattice-Boltzmann-immersed-boundary-finite-element solver to model near-rigid particles flowing through a straight microchannel with a rectangular cross-section. The simulation setup replicates the device introduced by Zhou et al. [12], and susequently used by Macaraniag et al. The device features three equally sized inlets and outlets. The two outer inlets introduce a blood sample containing white blood cells (WBCs) and circulating tumour cells (CTCs), while the inner inlet contains a buffer fluid with matching fluid properties. The intended outcome is for CTCs to exit through the central outlet and WBCs to exit through the outer outlets.

We first simulate single-particle behaviour for a range of particle sizes to establish a baseline behaviour and isolate the particle-particle interaction effects. We characterise particle trajectories, identify confinement-dependent heteroclinic orbits, and quantify migration velocities along these orbits. We then simulate a dilute suspension representative of experimental conditions to investigate the influence of CTCs on the dynamics of WBCs at both macroscopic and microscopic levels.

We demonstrate that particle migration is governed by size-dependent heteroclinic orbits. Larger particles, such as CTCs, tend to migrate closer to the channel centerline, while smaller WBCs remain near the walls. Importantly, migration velocities differ along these orbits, enabling velocity-based separation. However, this idealised behaviour is significantly altered in the presence of multiple particles.

Our results show that the presence of a single CTC can substantially influence the migration dynamics of surrounding WBCs, accelerating their lateral migration even across the channel centerline. These particle-particle interactions can lead to unexpected WBC presence in the target outlet, reducing separation purity. Our data confirm that such effects are most pronounced at short axial distances and in regions of higher local particle concentrations.

These findings highlight the limitations of single-particle models in predicting device performance under realistic conditions. To maintain high separation fidelity, IMF devices must operate under sufficiently dilute conditions to minimise hydrodynamic interactions. Furthermore, careful control of initial particle positioning and channel geometry may improve separation efficiency by exploiting velocity-dependent migration behaviours.

Ultimately, this work highlights the necessity of incorporating multi-particle dynamics into the design and analysis of microfluidic separation devices, contributing to more reliable CTC detection systems for early cancer diagnostics.

## Appendix

**Figure A1:**
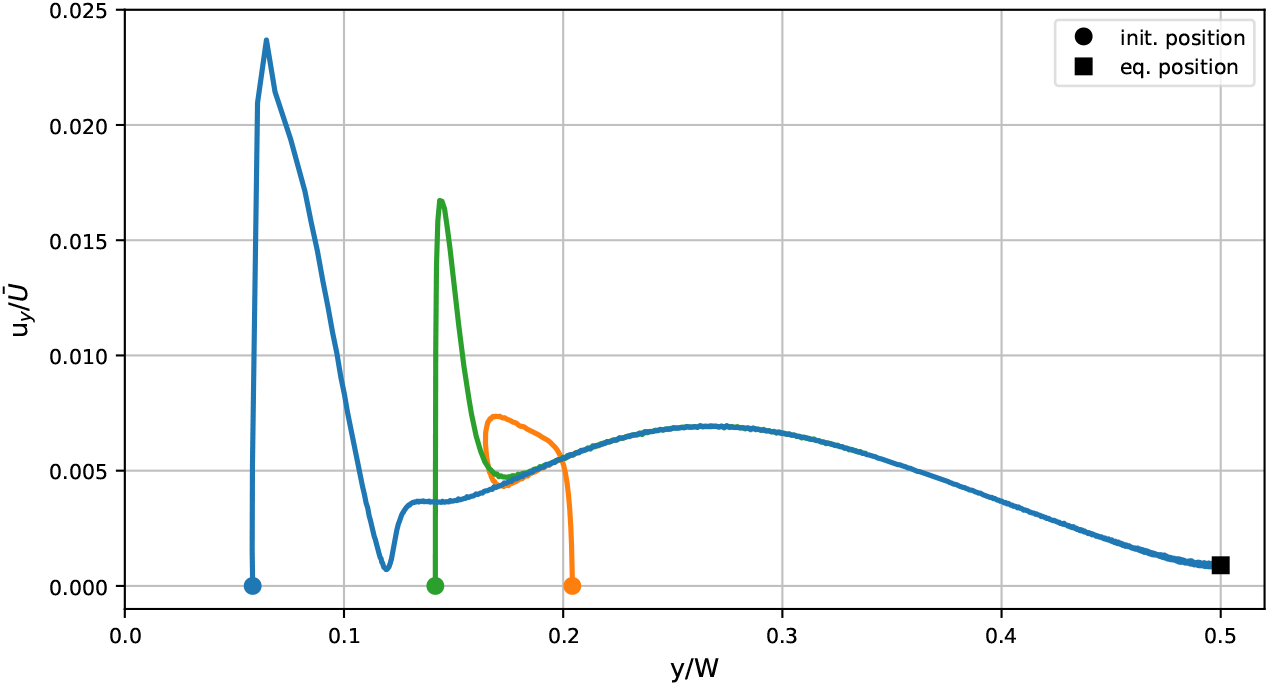
Lateral migration velocity along width direction as function of lateral position for three different initial positions of the same particle, *χ* = 0.34. Once the particle is on the heteroclinic orbit (*y/W >* 0.2), the lateral velocity only depends on particle location.

